# Validation of marker-less pose estimation for 3D kinematics during upper limb reaching

**DOI:** 10.1101/2023.03.16.532711

**Authors:** Inbar Avni, Lior Landau, Galya Shaked, Anat Shkedy Rabani, Raziel Riemer, Ahmet Arac, Lior Shmuelof

**Affiliations:** Department of Cognitive and Brain Sciences, Ben Gurion University of the Negev, Beer Sheva, Israel; Department of Industrial Engineering and Management, Ben Gurion University of the Negev, Beer Sheva, Israel; Department of Neurology, David Geffen School of Medicine, UCLA, Los Angeles, CA, USA

## Abstract

Kinematic analysis of movement following brain damage is key for diagnosing motor impairments and for recovery assessment. Advances in computer vision offer novel marker-less tracking tools that could be implemented in the clinic due to their simple operation and affordability. An important question that arises is whether marker-less technologies are sufficiently accurate compared to well established marker-based technologies. This study aims to perform validation of kinematic assessment using two high-speed cameras and a 3D pose estimation model. Four participants performed reaching movements with the upper limb between fixed targets, in different velocities. Movement kinematics were simultaneously measured using the DeepBehavior model and marker-based optical motion capture (QTM), as a gold standard. The differences in corresponding joint angles, estimated from the two different methods throughout the analysis, are presented as a mean absolute error (MAE) of the elbow angle. Quantitatively, the MAE of all movements was relatively small across velocity and joints (~2°). In a condition where the movements were made towards the DeepBehavior cameras, and the view of the elbow was occluded in one of the cameras, the errors were higher. In conclusion, the results demonstrated that marker-less motion capture is a valid alternative to marker-based motion capture. Inaccuracies of the DeepBehavior system could be explained by occlusions of key-points and are not associated with failure of the pose estimation algorithm.

## Introduction

The study of movement kinematics has taken leaps and bounds in the past decades with the emergence of motion capture technologies. Advancements in biological motion tracking have been widely utilized in industrial applications, sport, medicine, and science [1]. The leading gold standard in the field is camera-based markers tracking technology, which has been proven as highly accurate and reliable [2].

While these tools allow for detailed kinematic analysis of behavior, they require extensive expertise, long preparation times of the subjects, dedicated spaces, and are expensive. These limitations have brought into existence other motion capture technologies, at the forefront of which stand the utilization of novel object detection deep learning algorithms. These marker-less tools are trained to detect key points of human pose in 2D images, without the need for markers or specialized equipment [3–9]. Combined with transformation methods of 2D data of multiple cameras and depth estimation, these models can provide 3D kinematics of human motion [10–13].

To integrate these systems into research in motor control and in rehabilitation, their accuracy with respect to the gold-standard measurements should be determined. This research has been started, focusing primarily on gait analysis [14–17] or full body tasks [11]. The accuracy of marker-less pose estimation for upper extremity functions, which are frequently impaired following brain damage, has not been established. The aim of this study is to validate kinematic assessment of upper limb reaching movements using two high-speed cameras and a 3D pose estimation model (DeepBehavior). The validation will focus on the angles of the elbow and the shoulder during movements performed perpendicularly and in parallel to the DeepBehavior cameras, at three speeds. This validation is an essential step towards integration of marker-less pose estimation in the clinical environment for studying and assessing motor performance after neural and peripheral damage.

## Materials and methods

### Participants

Five healthy college-aged subjects participated in the study (4 females, age mean and std: 28.1±1.49). The study was approved by Ben-Gurion University’s internal reviewing board (#1317) and participants signed the approved consent forms. Recordings of one participant were excluded due to issues with Qualisys cameras being turned off during the recording session.

Experimental design: Each subject was seated and asked to perform reaching movements with their right arm to fixed targets positioned to their right side, or to their front (Fig. 1a-b). Targets were positioned 22 cm or 35.8 cm horizontally and 47.5 cm or 58.2 cm vertically from the origin point, depending on the task. Duration was controlled using a metronome, dictating movements at three durations: 40, 60 or 80 beats per minute (BPM). Each reaching condition was repeated 30 times in each duration.

**Figure 1.**
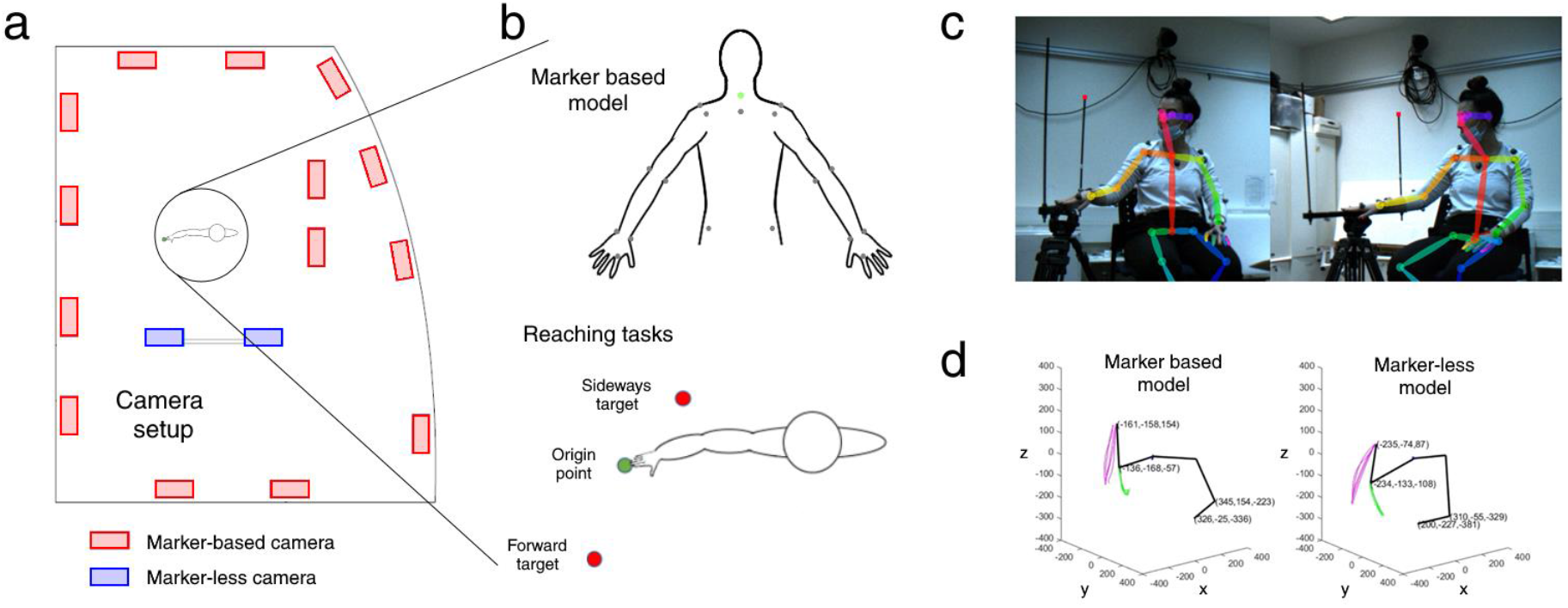
Experiment paradigm. (a) Camera setup of the two systems. (b) Marker locations and target location diagram. (c) Example frames from the two-camera system depicting the OpenPose model of one of the subjects sitting in the starting position. (d) The upper limbs of the 3D model of the marker-based system (QTM) and the marker-less system (DeepBehavior). Shoulder, elbow and wrist trajectories are plotted in blue, green and magenta respectively.

Data recording: Movements were recorded simultaneously using (1) a two-camera system for DeepBehavior analysis, and (2) multi-camera marker-based motion capture system (Qualisys, 150 Hz) for a gold standard analysis (Fig. 1a). In addition, we collected data with a low-tech phone camera (30 Hz) and an IMU accelerometer. We report here the results of the comparison between the DeepBehavior and the Qualisys systems.

### Marker-less motion capture

As previously reported [10, 18], the tasks were recorded using a custom-made system comprising of two high-speed cameras (150 frames per second, 1280×1024 pixels, Blackfly S Color 1.3 MP USB3 camera (FLIR Inc) with a Fujinon 1.5MP 6mm C Mount lens), set on a custom-designed aluminum camera holder with a 66° angle between their axes. Participants were recorded from the frontal angle while facing the cameras, set 2.2 m away from them and at the level of shoulders when the subjects were sitting. The data from the two cameras allowed us to produce the 3D DeepBehavior kinematic model consisting of 57 body joints (Fig. 1c-d).

### Marker-based motion capture

Before each session, subjects were fitted with a set of 18 markers on the arms and torso: 7 markers on each arm (2 on the shoulder, 2 on the elbow, 2 on the wrist and another one on the first knuckle), 2 markers on the hips, and 2 markers on the chest and back (Fig. 1b). An infra-red tracking system composed of 14 motion capture cameras mounted on the walls surrounding the subject at an average height of 2.56 m (Qualisys Oqus 5+, 4Mpixel 180Hz) were used to detect the markers that are located on the subjects’ body (Fig. 1a). Calibration of the cameras was completed before each recording session. 3D positions of the markers were recorded in 150 Hz using the Qualisys Track Manager recording software.

### Data analysis

The marker-less 3D model kinematic data was extracted from the raw videos of the two-camera system. Each camera footage was processed using the OpenPose toolbox [7]. The two estimates were then stereo-triangulated to obtain the 3D positions of the joints using the DeepBehavior toolbox [10]. Marker-based 3D data was produced using a customized software of the system: Qualisys Track Manager (QTM) for labeling and extracting the 3D position of the markers. Data analysis was performed using a custom written code in MATLAB [19].

Both systems data sets were smoothed using a Savitzky-Golay filter with a window size of 31 and a polynomial degree of 3. Then the two models were synchronized for each task recording using the alignsignals Matlab function. Movements were segmented in the two models based on the tangential velocity profiles of the wrist marker position of the right arm in the marker-based model. Start and end points of individual movements were defined based on the crossing of one standard deviation below the peak velocity mean, before and after the peak velocity, respectively. Based on the segmentation of the marker-based 3D data, the marker-less 3D data was segmented.

For the marker-based system, joint position estimation for the shoulder and elbow joints was based on the average of the two markers positioned on each joint, while the position of the wrist was set as the marker placed on the internal-lateral aspect of the wrist (figure 1b). For the marker-less system, joint position estimation was based on the position of the shoulder, elbow and wrist extracted from the DeepBehavior model.

Joint angle data was calculated in intrinsic (anatomical) coordinate system. Angles were calculated in relation to a specific joint: shoulder flexion angle was defined as the angle in degrees between the ipsilateral elbow joint, ipsilateral shoulder joint and contralateral shoulder joint, projected onto the horizontal plane – defined by the torso and shoulder vectors (defined from the chest point to the midhip point and from the right shoulder to the left shoulder, respectively). Elbow extension angle was defined as the angle in degrees between the ipsilateral wrist, elbow and shoulder joints.

After joint angles were calculated for the movement trajectories, angle trajectories of the marker-based system were normalized so the angle value at the end of the movement is zero. The angle trajectories of the marker-less system were then aligned to the marker-based data using a minimum MAE criterion.

### Statistical analysis

In order to study the accuracy of the marker-less system, we calculated the mean absolute error (MAE) between the systems by computing the absolute difference between the joint angles in each time-point along the movement and averaging across them. ANOVA was then used for detecting biases between systems. Pearson correlation was calculated between the angle trajectories of the two systems to characterize the correspondence between the systems.

## Results

### Validity of the model in different velocities and different joints

The elbow angle trajectories in different velocities are presented in figure 2. Results of the two models are very similar for all subjects with very high correlation coefficient values and low mean absolute errors (mean and std across participants and velocities 2.01°±0.82°; maximum error 3.2°). No significant effect of velocity on MAE was observed.

**Figure 2.**
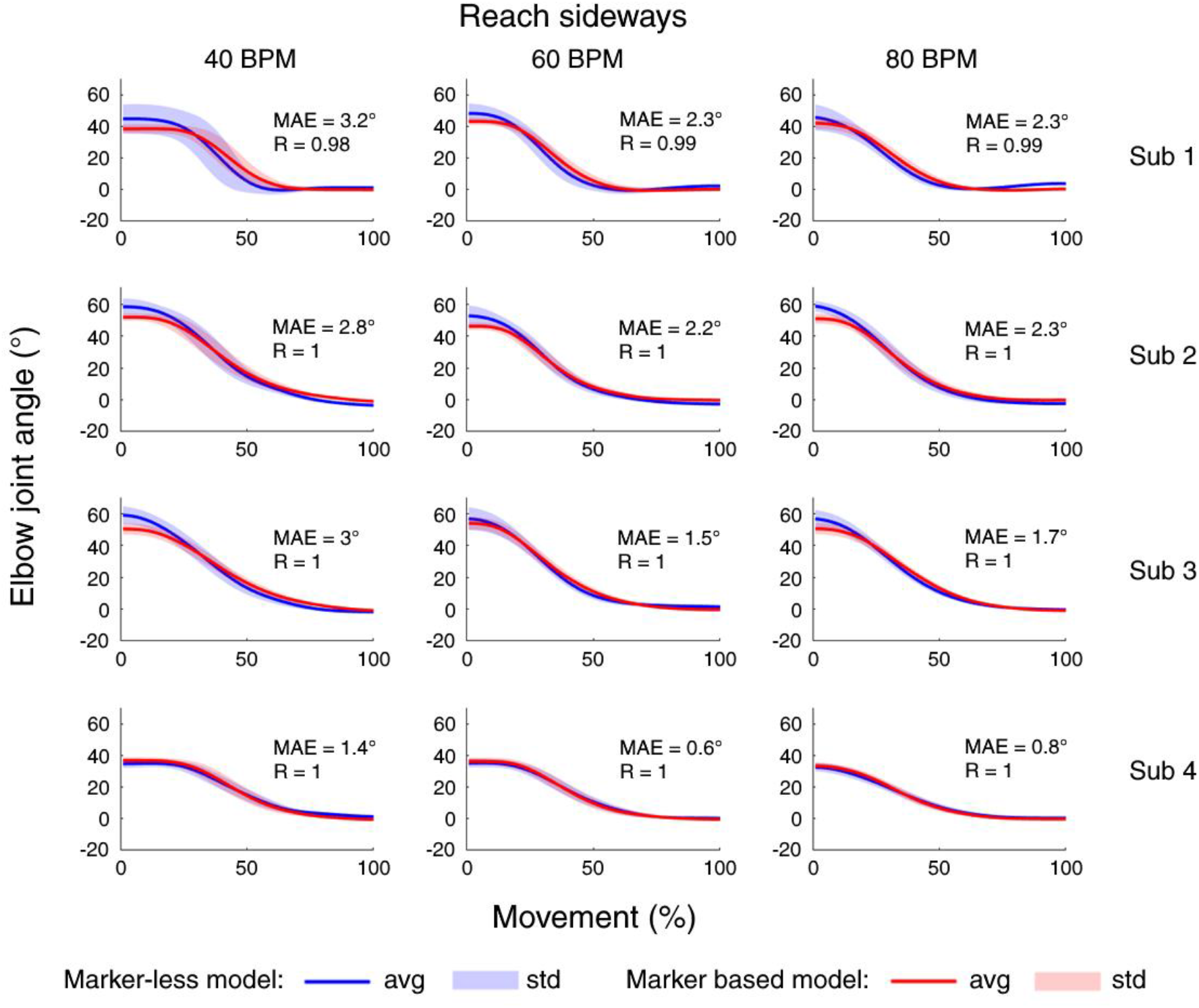
Comparison of elbow angle changes during movement in different velocities. Elbow angle averaged across individual movements in the marker-less DeepBehavior model (blue) and marker-based QTM model (red). Shaded area represents the standard deviation of each data set. Each row represents a single subject and each column represents a different velocity (40 BPM, 60 BPM and 80 BPM). Mean average error and Pearson correlation values calculated between the two systems are noted.

Shoulder angle results compared to the elbow angle are presented in figure 3. Mean absolute errors for the shoulder are also relatively low in most subjects (mean and std across participants: 2.O2°±1.6°; maximum error 4.3°). No significant effect of joint on MAE was observed.

**Figure 3.**
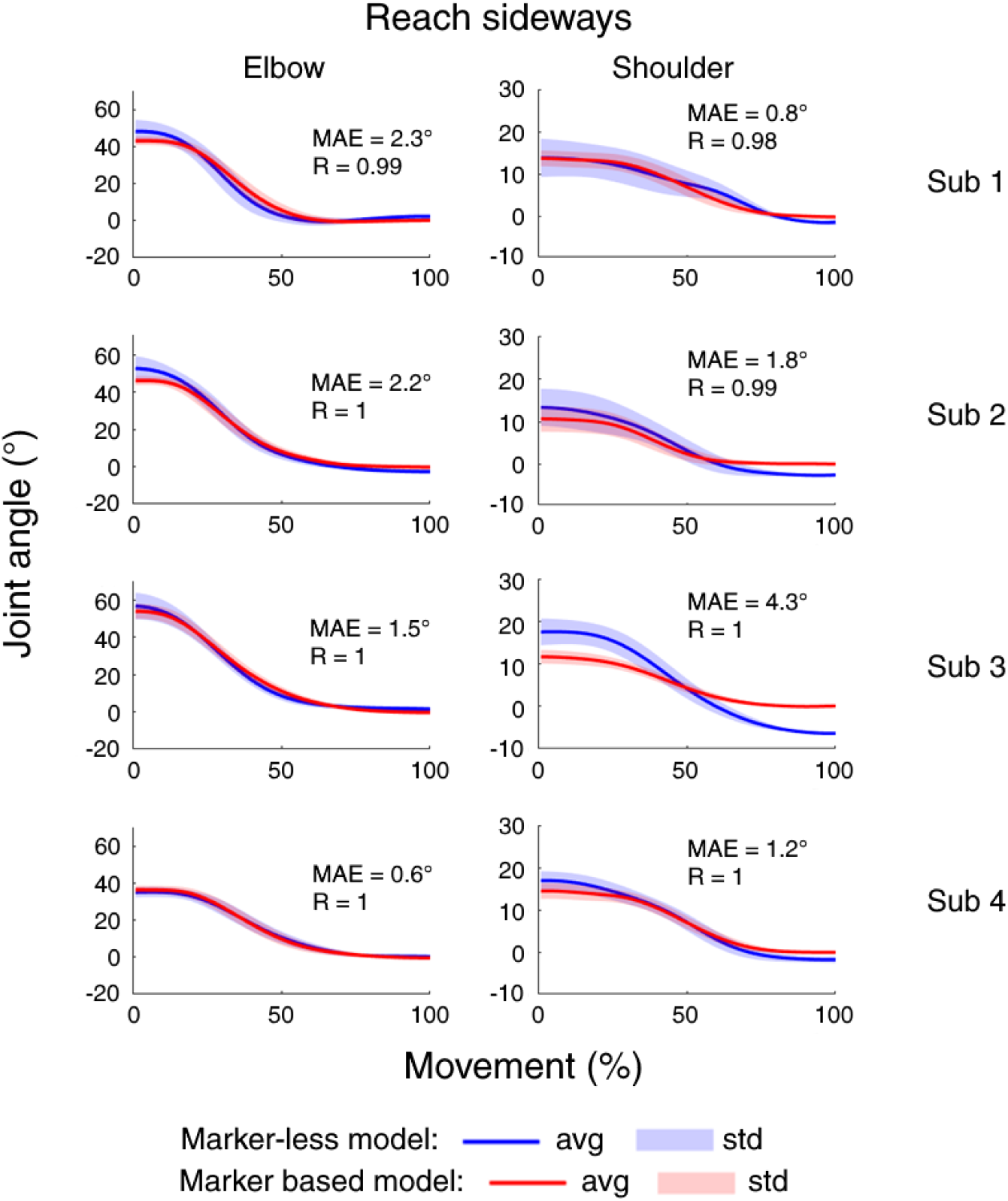
Comparisons of elbow and shoulder angle changes during movement. Elbow and shoulder angles averaged across individual movements in the marker-less DeepBehavior model (blue) and marker-based QTM model (red). Shaded area represents the standard deviation of each data set. Each row represents a single subject. Mean average error and Pearson correlation values calculated between the two systems are noted.

### Validity in conditions of occlusion

After assessing the accuracy of the DeepBehavior system in conditions where the movements were parallel to the cameras, we wanted to assess the accuracy of the system in conditions where the movement is perpendicular to the main axis of one of the cameras. To do so, we looked at the accuracy of the marker-less model in a task consisting of movement to a target presented in front of the subject. Indeed, the deviation of the DeepBehavior system increased (Fig 4), showing larger MAEs for all subjects (mean and std across participants and velocities 7.13°±5.2°; maximum error 12.2°).

**Figure 4.**
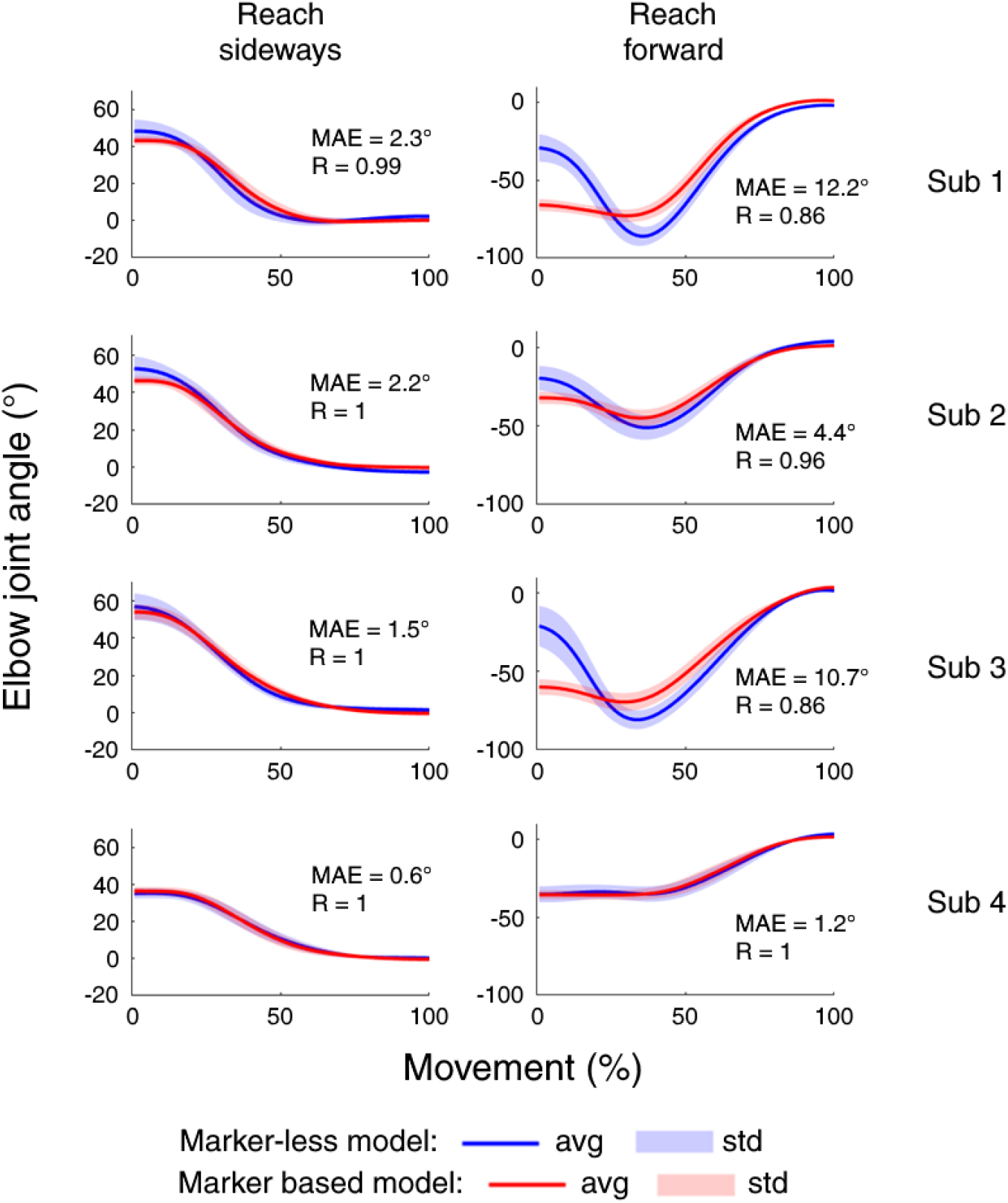
Elbow angle trajectory mean in conditions of occlusion. Elbow angle averaged across individual movements in the marker-less DeepBehavior model (blue) and marker-based QTM model (red). Shaded area represents the standard deviation of each data set. Each row represents a single subject. The angle trajectories are presented for the movement towards the target set to the side (left) and the target set to the front of the subject which included occlusion (right). Mean average error and Pearson correlation values calculated between the two systems are noted.

To examine if these inaccuracies are driven by occlusions, we calculated the distance between the wrist and elbow joints of the OpenPose 2D model from each camera and compared it to the level of inaccuracy between the models of the elbow angle, in the forward target task. As expected, recordings that had occlusions (defined as less than 75 pixels (3.36°) between wrist and elbow, see figure 5), showed inaccuracies in elbow angle estimation (figure 6). Furthermore, for the subject that showed no occlusions, the accuracy of the DeepBehavior systems was the highest, with a MAE of 1.2° (subject 4 in figure 6a).

**Figure 5.**
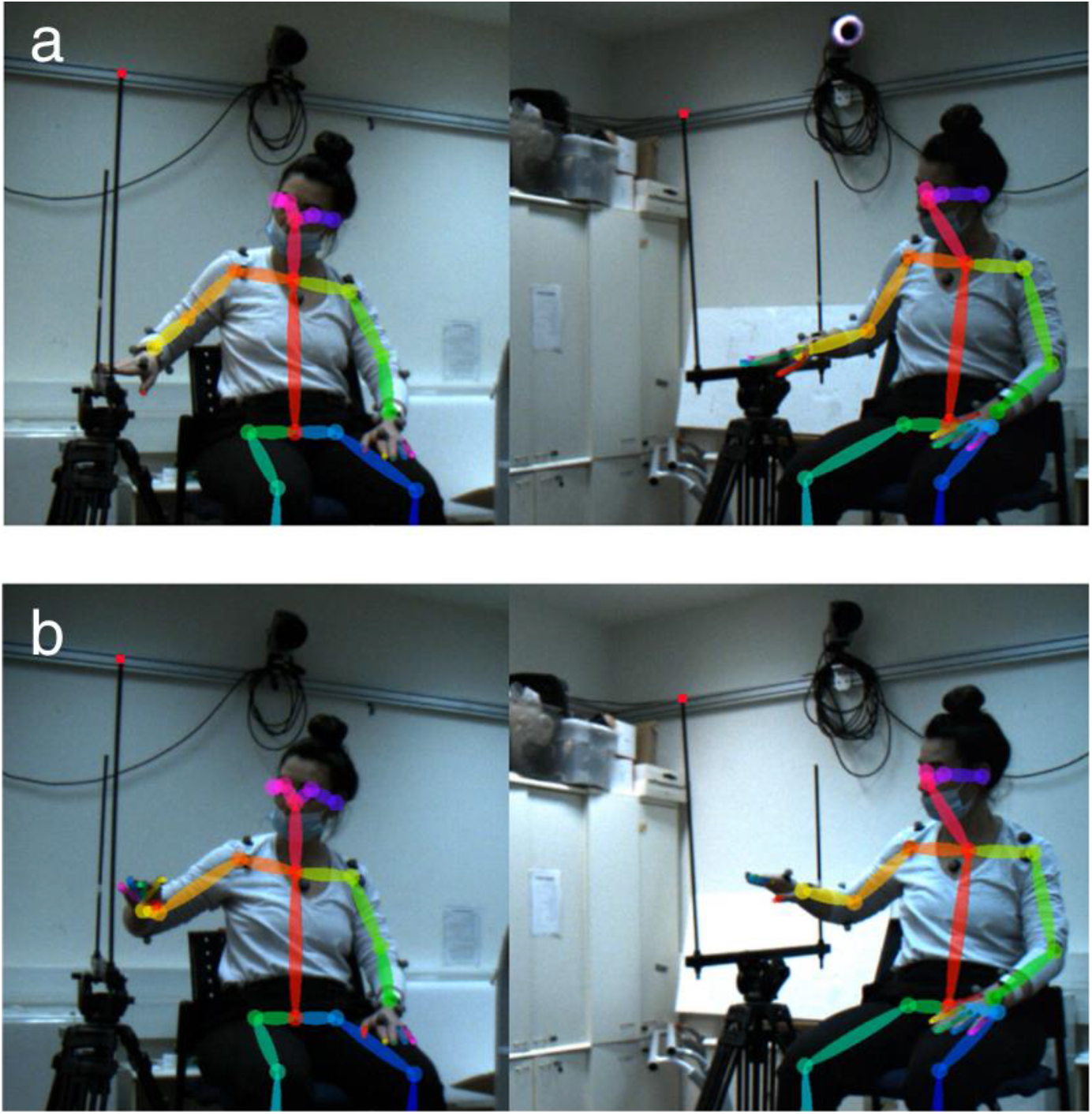
Example of elbow occlusion in one of the subjects performing reaching for the forward target. The 2D OpenPose model for a single subject performing reaching for the forward target from camera 1 (left) and camera 2 (right), at the start **(a)** and middle of the movement **(b**). The occlusion is in panel b, picture on left (camera 1) (the right elbow and wrist of the subject are in close proximity in the camera view).

**Figure 6.**
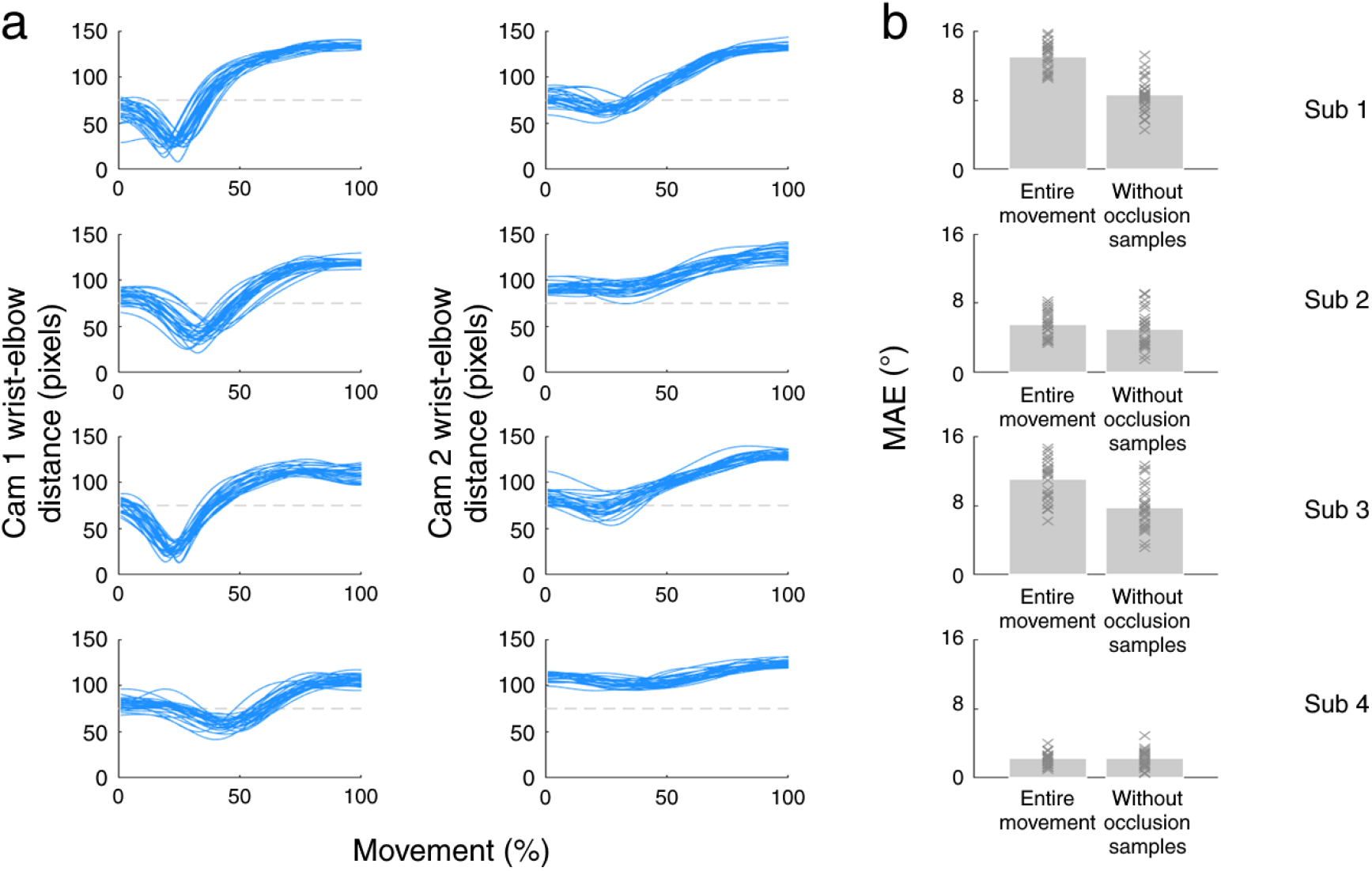
Model accuracy in conditions of occlusions. Distance between wrist and elbow joints positions in the 2D model of OpenPose in camera 1 (left) and camera 2 (middle) (**a**). Occlusion threshold of 75 pixels is noted in gray dashed horizontal line. Mean absolute error (MAE) of angle trajectories of the two 3D models for each subject performing reaching for the forward target (**b**). MAE is presented for individual trials (noted as x in dark gray) and averaged across trials (bars in light gray), for the entire trial and for the entire trial excluding the samples which had occlusion over a threshold of 75 pixels.

To quantify the effect of occlusion on accuracy, we compared the of MAE the entire trial with the MAE of the trial without the frames with occlusion (i.e., frames where the distance between the wrist and elbow joints of the 2D model, from at least one of the cameras, was smaller than 75 pixels). The MAE for trajectories without occlusion was smaller for all subjects (mean and std across participants 5.87°±2.88°; figure 6b).

## Discussion

This study aimed to examine the accuracy of the OpenPose-based 3D marker-less motion capture model DeepBehavior, through a comparison with an optical marker-based motion capture (Qualisys). From the data presented here, the DeepBehavior model produces kinematic variables that are comparable across subjects, velocities (figure 2), and joints (figure 3).

We calculated the MAE of the elbow angle during movement. Overall, the errors are around 2 degrees and the correlation coefficients of individual subjects across tasks and velocities are close to one.

To test the accuracy of the system in sub-optimal conditions, we examined its accuracy in a task that required subjects to move the arm towards the cameras. We found that that accuracy of the system is sensitive to such a manipulation, and that inaccuracies can be explained by occlusions (figures 4–6). This has been previously shown as a limitation for a two-camera system, that can be solved by either using additional cameras or adjusting the angle of the cameras with respect to the movement of the subject [20]. In any case, it suggests that inaccuracies are not driven by failure of the pose estimation models, but by the sub-optimal input to the DeepBehavior model.

The results demonstrate that marker-less motion capture is a valid alternative to marker-based motion capture when used carefully. This result paves the road towards integrating similar systems in clinics and developing kinematic measures for diagnosing motor impairment and assessing recovery and response to treatment. A recent round table discussion emphasized the importance of kinematic movement quantification for assessing stroke recovery [21]. We propose that marker-less kinematics is a feasible and valid approach to achieve that goal.

### Limitations

This study validated important kinematic measures of upper extremity; however, additional validation is needed for estimating the position of the fingers and of the wrist, that are typically noisier and more susceptible to occlusions. Additionally, pose estimation in subjects with neural and motor disorders may suffer from inaccuracies due to the untypical postures presented by these subjects when compared to the training data set. Thus, validation in specific clinical groups may be also needed.

### Conclusions

We show that an affordable and mobile two-camera system, combined with a pose estimation algorithm, can provide accurate pose estimation data and 3D kinematic measures that are highly correlated with the results of a gold standard marker-based pose-estimation system. Such systems can be positioned in clinics and support diagnosis of motor impairments and recovery assessments following injury, and for scaling the research of motor pathologies using kinematics.

